# A 2.08 Å resolution structure of HLB5, a novel cellulase from the anaerobic gut bacterium *Parabacteroides johnsonii DSM 18315*

**DOI:** 10.1101/516542

**Authors:** Changsoo Chang, Charles Brooke, Hailan Piao, Jamey Mack, Gyorgy Babnigg, Andrzej Joachimiak, Matthias Hess

## Abstract

Cellulases play a significant role in the degradation of complex carbohydrates. In the human gut anaerobic bacteria are essential to the well-being of the host by producing these essential enzymes that convert plant polymers into simple sugars that can then be further metabolized by the host. Here we report the 2.08 Å resolution structure of HLB5, a chemically verified cellulase that was identified previously from an anaerobic gut bacterium and that has no structural cellulase homologues in PDB nor possesses any conserved region typical for enzymes that degrade carbohydrates. We anticipate that the information presented here will facilitate the identification of additional cellulases for which no homologues have been identified until to date and in enhancing our understanding how these novel cellulases bind and hydrolyze their substrates.

## Introduction

Plant carbohydrates composed of large polymers such as cellulose, hemi-cellulose, and lignin are a major energy source for microorganisms that inhabit the digestive tract of animals and humans. To efficiently decompose carbohydrate polymers these microbes utilize a diverse set of carbohydrate active enzymes (CAZymes; www.cazy.org) (Lombard et al. 2014). Although in some bacterial genomes CAZymes account for more than 10% of the encoded genes, it is believed that many are still to be discovered. Cellulose, a polymer of glucose, represents a major fraction of plant carbohydrates and three classes of cellulases are needed to decompose cellulose: 1,4-β-endoglucanases, β-glucosidases. Among these, 1,4-β-endoglucanases (endoglucanase; EC 3.2.1.4), randomly hydrolyze 1,4-β-glucosidic bonds, represent a key group in the degradation of cellulose (Lynd et al. 2002). Besides their physiological importance, CAZymes have recently received increased attention due to their industrial potential (Dashtban et al. 2010; Lynd et al. 2017; Kuhad et al. 2011; Harris and DeBolt 2010; Tan et al. 2019).

In a previous study we utilized a “guilty by association” strategy to discover novel cellulases (Piao et al. 2014). We identified and cloned 17 putative cellulases with too little sequence similarity to known cellulases to be identified as such. From this set, 11 (~65%) were verified to possess cellulolytic activity against carboxymethylcellulose (CMC) and pretreated *Miscanthus* (Piao et al. 2014). Of these cellulases, HLB5 possessed high activity towards both CMC and *Miscanthus*. In the genome of *Parabacteroides johnsonii* DSM 18315, *hlb5* is part of a gene cluster, containing among others a β-galactosidase/glucuronidase, two β-1,4-xylanases, a sugar phosphate permease, and three bacterial outer membrane proteins. Further analysis of the 247 amino acid long sequence of HLB5 failed to detect any signal peptide or transmembrane helices but identified a region (PF03737) conserved in proteins belonging to the RraA and RraA-like family.

Here, we present the crystal structure of HLB5 at 2.08 Å resolution. The protein has no significant sequence identity to known cellulases. The structure shows tight hexamer and is providing information about the three-dimensional arrangements and functional sites. HLB5 has several structural homologues in PDB, none of them have cellulase activity and the protein does not possess any conserved regions typical for CAZymes whilst still being capable of degrading cellulosic biomass.

## Materials and methods

### Sub-cloning, expression and purification

Residues 3 to 243 of *hlb5*, was amplified by PCR from the previously constructed *p*ET102-*hlb5* (Piao et al. 2014) with KOD Hot Start DNA polymerase (MilliporeSigma, Hayward CA, USA) with forward (5’-TACTTCCAATCCAATGCCGTAGATCAGTATAAGAAAGAAATCGGAATGATGA-3’) and reverse primers (5’-TTATCCACTTCCAATGTTATTTTGCTGTAATTTCCTGAACACTGTCTC-3’). The PCR product was purified and cloned into pMCSG68 (MCSG, Argonne, IL, USA) using modified ligase-independent procedure (Stols et al. 2002) and transformed into *Escherichia coli* BL21(DE3)-Gold strain (MilliporeSigma, Hayward CA, USA). The pMCSG68 vector provides an N-terminal TEV-cleavable His6 purification affinity tag. After cloning and sequencing of the insert, a point mutation of Pro191 to Ser was detected. Cells were grown in enriched M9 medium using selenomethionine (SeMet), under conditions known to inhibit methionine biosynthesis at 37°C to an OD_600_ of 1. Protein expression was induced with 0.5 mM IPTG. Expression was conducted overnight under aeration (200 rpm) at 18°C. Cells were harvested by centrifugation (2,200 × *g*) and resuspended in five volumes of lysis buffer (50 mM HEPES pH 8.0, 500 mM NaCl, 20 mM imidazole, 10 mM 2-mercaptoethanol and 5% v/v glycerol) and stored at −20°C until processed further. Harvested cells were thawed and subsequently treated with a protease inhibitor cocktail (P8849, MilliporeSigma, Hayward CA, USA) and 1 mg/ml lysozyme prior to sonication. The lysate was clarified by centrifugation at 30,000 × *g* (Sorvall RC5C-Plus, ThermoFisher, West Sacramento, CA, USA) for 60 min, followed by filtration through 0.45 and 0.22 μm in-line filters (Gelman, Pall Corporation, Westborough, MA, USA). The protein was purified with immobilized metal affinity chromatography (IMAC-I) using a 5-ml HiTrap Chelating HP column charged with Ni^2+^ ions followed by buffer-exchange chromatography on a HiPrep 26/10 desalting column (both GE Healthcare Life Sciences, Pittsburgh, PA) using an ÄKTAxpress system (GE Healthcare Life Sciences). The His_6_-tag was cleaved using the recombinant TEV protease expressed from the vector pRK508 (MCSG, Argonne, IL). The protease was added to the target protein in a ratio of 1:30 and the mixture was incubated at 4°C for 48 h. The protein was then purified using a 5 ml HiTrap Chelating column charged with Ni^2+^ ions. The protein was dialyzed in 20 mM pH 8.0 HEPES, 250 mM NaCl, 2 mM DTT and concentrated using a Centricon Plus-20 Centrifugal Concentrator (MilliporeSigma, Hayward, CA, USA) to 57 mg/ml.

### Protein crystallization and data collection

Crystallization conditions were determined using MCSG crystallization suite (Mycrolytic) with the help of Mosquito robot (TTP Labtech) using sitting drop vapor diffusion technique in a 96-well CrystalQuick plate (Greiner). Crystals suitable for X-ray diffraction data collection were grown in a condition of MCSG3 well 43 (0.2M Li_2_So_4_, 0.1 M CHES pH9.5, 0.1M Na/K Tartrate) at 24°C. Single wavelength anomalous diffraction data near the selenium absorption peak was collected from a SeMet-substituted protein. The crystal growth buffer was supplemented with 25% ethylene glycol for cryoprotectantion. The crystal was picked using a Litholoop (Molecular dimensions, Maumee, OH, USA) and flash-cooled in liquid nitrogen. Data were collected on an ADSC quantum Q315r charged coupled device detector (Poway, CA, USA) at 100°K in the 19ID beamline of the Structural Biology Center at the Advanced Photon Source, Argonne National Laboratory. The crystal belongs to hexagonal space group P6_3_22 with cell parameters of *a*=*b*=100.2 Å, *c*=129.7 Å, α=β=90°, γ=120°. The diffraction data were processed by the HKL3000 suite of programs (Minor et al. 2006). Data collection statistics are presented in Table 1.

**Table 1.** Data collection and refinement statistics.

|  |  |  |
| --- | --- | --- |
| Data collection statistics |  |  |
|  | Wavelength (Å) | 0.97921 |
|  | Resolution (Å) | 50-2.08 |
|  | Space group | P6<sub>3</sub>22 |
|  | Unique reflections | 23,828 |
|  | Completeness (%) | 100(100) |
|  | I/sigma | 39.1(3.09) |
|  | R<sub>merge</sub> (%) | 0.11(0.98) |
| | Cell parameters (Å, °) | $a = b = 100.2$, $c = 129.7$,<br>$\alpha = \beta = 90$, $\gamma = 120$ |
| Refinement statistics |  |  |
|  | Resolution range (Å) | 41.2-2.08 |
| | $R_{\text{work}}/R_{\text{free}}$ (%) | 14.2/18.7 |
|  | Number of |  |
|  | atoms | 1996 |
|  | protein atoms | 1767 |
|  | hetero atoms | 31 |
|  | water atoms | 198 |
| Root mean square deviation |  |  |
|  | Bond lengths (Å) | 0.002 |
|  | Angles (°) | 0.47 |
| Ramachandran plot |  |  |
|  | Favored | 98.20% |
|  | Outliers | 0% |
| Average B factor |  |  |
|  | All atoms (Å<sup>2</sup>) | 35.5 |
|  | Protein atoms (Å<sup>2</sup>) | 34.4 |
|  | Ligand atoms (Å<sup>2</sup>) | 42.6 |
|  | Solvent atoms (Å<sup>2</sup>) | 45.2 |

### Structure determination and refinement

All procedures for single wavelength anomalous diffraction (SAD) phasing, phase improvement by density modification, and initial protein model building were done using the structure module of the HKL3000 software package (Minor et al. 2006). Seven selenium sites were found using SHELXD (Sheldrick 2010). The mean figure of merit of the phase set from MLPHARE (Otwinowski 1991) was 0.213 for 50–2.08 Å data and improved to 0.856 after density modification (DM) (Cowtan 1994). The structure building module using arp/warp (Langer et al. 2008) of the HKL3000 package built 199 out of 241 residues, while side chain of 187 residues were placed. The initial model was rebuilt manually with the program COOT (Emsley and Cowtan 2004) by using electron density maps based on DM-phased reflection file. After each cycle of rebuilding, the model was refined by PHENIX (Adams et al. 2010). The geometrical properties of the model were assessed by COOT and MolProbity (Davis et al. 2007).

### Structure determination and model quality

The crystal structure of HLB5 was solved by single wavelength anomalous diffraction method. There is one molecule in an asymmetric unit and the structure was refined to an R/R_free_ of 0.142/0.187 in 41.2-2.08 Å resolution range. The structure shows acceptable root mean square deviation from ideal geometry and reasonable clash score from Molprobity. Detailed refinement statistics are shown in Table 1.

## Results

### Structure of HBL5

The monomer structure is α/β/β/α sandwich-fold composed of 8 helices and 13 strands (Fig. 1A). The first layer is composed of α-helices α5, α6, α7 and small β-sheet, made of short β-strands β4, β6, β8. These small β-strands are usually composed of two amino acids. The second layer has a β-sheet composed of β-strands in the following order: β1, β7, β5, β3, β2, β9 The central four β-strands are in parallel, while the outside strands β1 and β9 are anti-parallel. Strand β2 is 12 amino acids long and bends to the third layer where it is involved in the β-sheet of the third layer. The third layer is composed of five β-stands that are forming two small β-sheets. β-strands β2, β12, and β13 make an anti-parallel β-sheet and strands β11 and β10 make the antiparallel β-sheet. The fourth layer is composed of five α-helixes (i.e. α1, α2, α3, α8, α4). Among these helixes, the first two helixes protrude toward an adjacent molecule to make contact.

**Figure 1.**
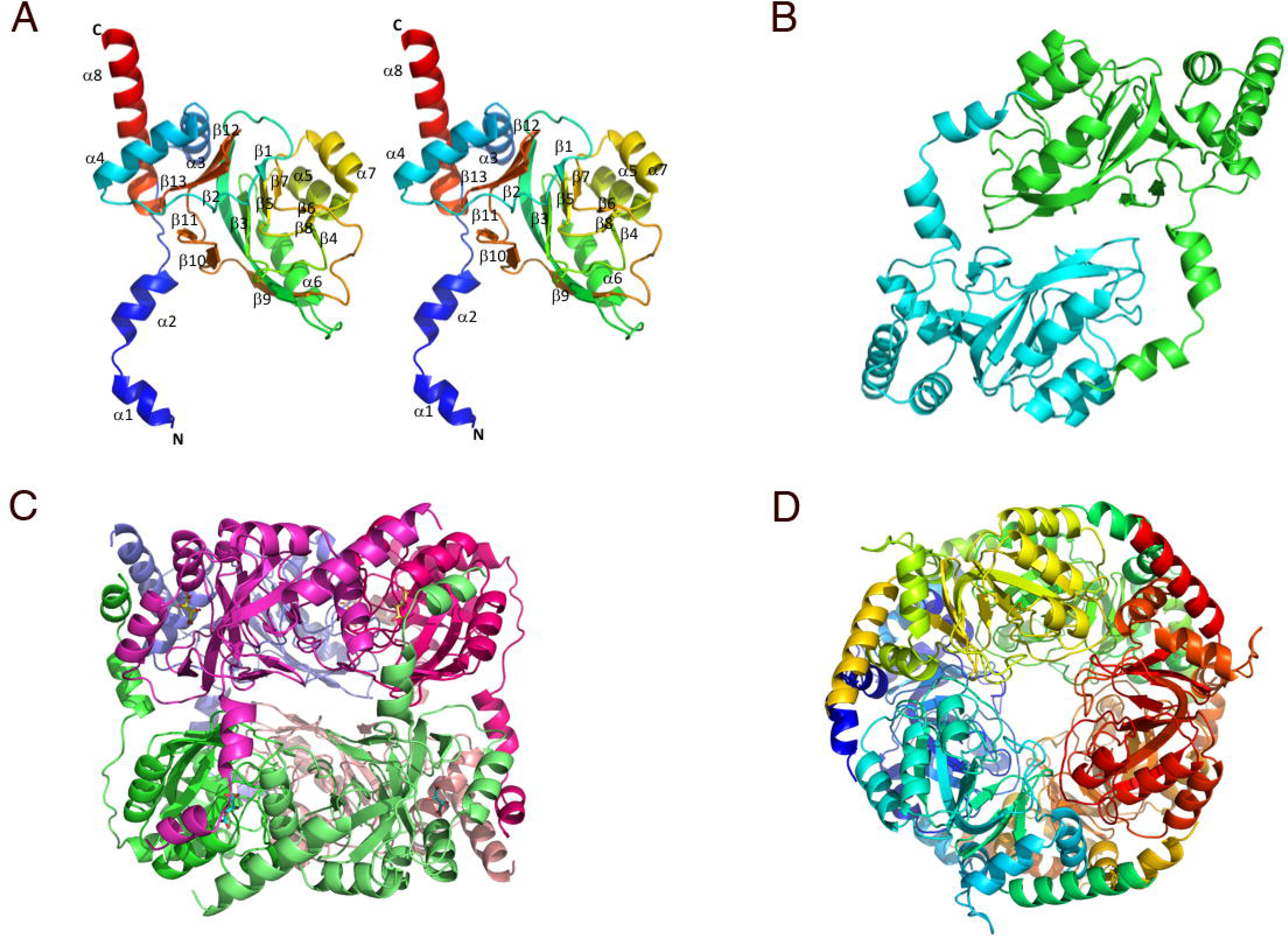
Crystal structure of HLB5. **(A)** A stereo view of the monomer structure. It is colored blue to red, from N-terminus to C-terminus. Secondary structure elements are labeled. **(B)** Dimeric assembly of HLB5, two monomers are colored green and cyan. **(C)** Hexamer of HLB5, view along 2-fold axis. Each chain is represented in different color. **(D)** Hexamer of HLB5, view along 3-fold axis, same color scheme as in C.

The oligomeric state of this protein was determined to be a hexamer in solution (data not shown), using size exclusion chromatography. The analysis of packing with PDBe PISA (Krissinel and Henrick 2007) suggests that the biological unit as hexamer, generated by the symmetric operators (x,y,z), (x,x-y, 1/2-z), (1–x+y,1-x,z), (1-y,1-x,1/2-z), (1-y,x-y,z), and (1–x+y,y,1/2-z). The hexamer displays perpendicular 3-fold and 2-fold symmetry. Two monomers bind each other tightly with the second and third layers from one polypeptide chain and α1, α2 helixes from another (Fig. 1B). The dimeric interface area is 3,630 Å^2^, accounting for 30% of the monomer’s surface (12,300 Å^2^). Three of these dimers are arranged in a ring shape around a 3-fold axis to make a hexamer with overall dimensions ~60 Å × 80 Å. There are also extensive interactions between neighboring dimers involving several helices and loop regions. A buried area of the hexamer is 27,800 Å^2^, while the surface area of the hexamer is 49,600 Å^2^ indicating very extensive interactions in the hexamer suggesting that this hexamer is very stable. The formation of a hexamer gains 64.7 Kcal/mol calculated using PISA (Krissinel and Henrick 2007).

Inspection of the hexamer revealed several clefts and a large channel located at the interfaces between subunits. The significant feature of the hexamer is a channel ~60 Å long running across entire structure along three-fold axis with diameter around 6 Å (Fig. 1D). In the middle of hexamer this channel expands and connects to a large cavity that appears to have three wide connections to the protein surface that are lined with acidic residues. There are also three narrow channels. All these channels and cavities are strongly hydrated and show a number of ordered water molecules. The large clefts located near a two-fold axis on the interface of four peptide chains (Fig. 1C) have wide entrance about 15 Å wide with 19 Å long, and the depth is about 15 Å leading to the central channel. In addition to the aforementioned opening, a smaller cleft not related to rotation axis exists. It is on the concave side of the sheet in layer 2 and surrounded by helixes α1, α4, α7 and an entrance is slightly covered by helix α1 of an adjacent molecule. This cleft appears to include ligand binding site.

### Homologous Structures

When the structure of HLB5 was subjected to the Dali server (Holm and Laakso 2016), no significant similarity to known cellulases was detected. The uncharacterized protein PSPTO_3204 from *Pseudomonas syringae* pv. tomato str. DC3000 (PDB 3k4i) had the highest Z score (26.3) followed by RNA-processing inhibitor RnaA like protein YER010CP from yeast (PDB 2c5q), 4-Hydroxy-4-methyl-2-oxoglutarate/4-Carboxy-4-hydroxy-2-oxoadipate(HMG/CHA) aldolase from *Pseudomonas putida* (PDB 3noj), S-adenosylmethionine/2-dimethylmenaquinone methyltransferase from *Geobacillus kaustophilus* (PDB 2pcn), RNA-processing inhibitor RraA’s from *Streptomyces coelicolor* (PDB 5×15), *Thermus thermophilis* (PDb 1j3l), *E. coli* (2yjv, 1q5x), *Vibrio cholera* (1vi4), *P. aeruginosa* (3c8o), putative Methyltransferase from *Mycobacterium tuberculosis* (1nxj) with Z scores over 15. Among these, HMG/CHA aldolase had a Z-score of 22.9. Interestingly, this HMG/CHA aldolase also has α/β/β/α sandwich-fold and exists as a hexamer (Wang et al. 2010) and when compared to HLB5, structures show nearly identical fold with r.m.s.d.(~ 1.9 Å). Differences of these structures are observed in the N- and C-termini of these sequences. More specifically, the N-terminus of HLB5 is protruding to interact with an adjacent molecule to generate dimers while the N-terminus of HMG/CHA aldolase does not. The C-terminus of HMG/CHA aldolase has 35 more residues than HLB5 and it extends out to wrap around an adjacent protomer of the trimer.

### Ligand Binding Site

The catalytic site of HMG/CHA aldolase is located in the cleft between adjacent protomers of the trimer and equivalent clefts are found on the structure of HLB5 (Fig. 2A). As described previously, this cleft is located in the region partially covered by the N-terminal helix of an adjacent polypeptide and occupied by a tartrate molecule from the crystallization solution. This tartrate makes hydrogen bonds with Gly125, Val127, Mse128, and Ser169, and hydrophobically interacts with Thr122, Trp124, Gly126, Arg147, and Gly168 (Fig. 2B). In the HMG/CHA aldolase structure, pyruvate and magnesium ions are bound to the active site. A pyruvate interacts with a magnesium ion and the side chain of residues Arg123 (Arg147 in HBL5), while the backbone N of residues Asp102, Leu103, and Leu104 (Gly126, Val127, Met128), and the magnesium ion interacts with Asp124 (Asp148 in HBL5). This spatial environment is well conserved although the metal ion is not found in HBL5 and different ligands are present. Other than these, the residues like Gly125, Gly168 are spatially conserved in this cleft. Trials for crystallization of putative substrate bound protein have failed so far and the exact active site residues of HLB5 remain unknown.

**Figure 2.**
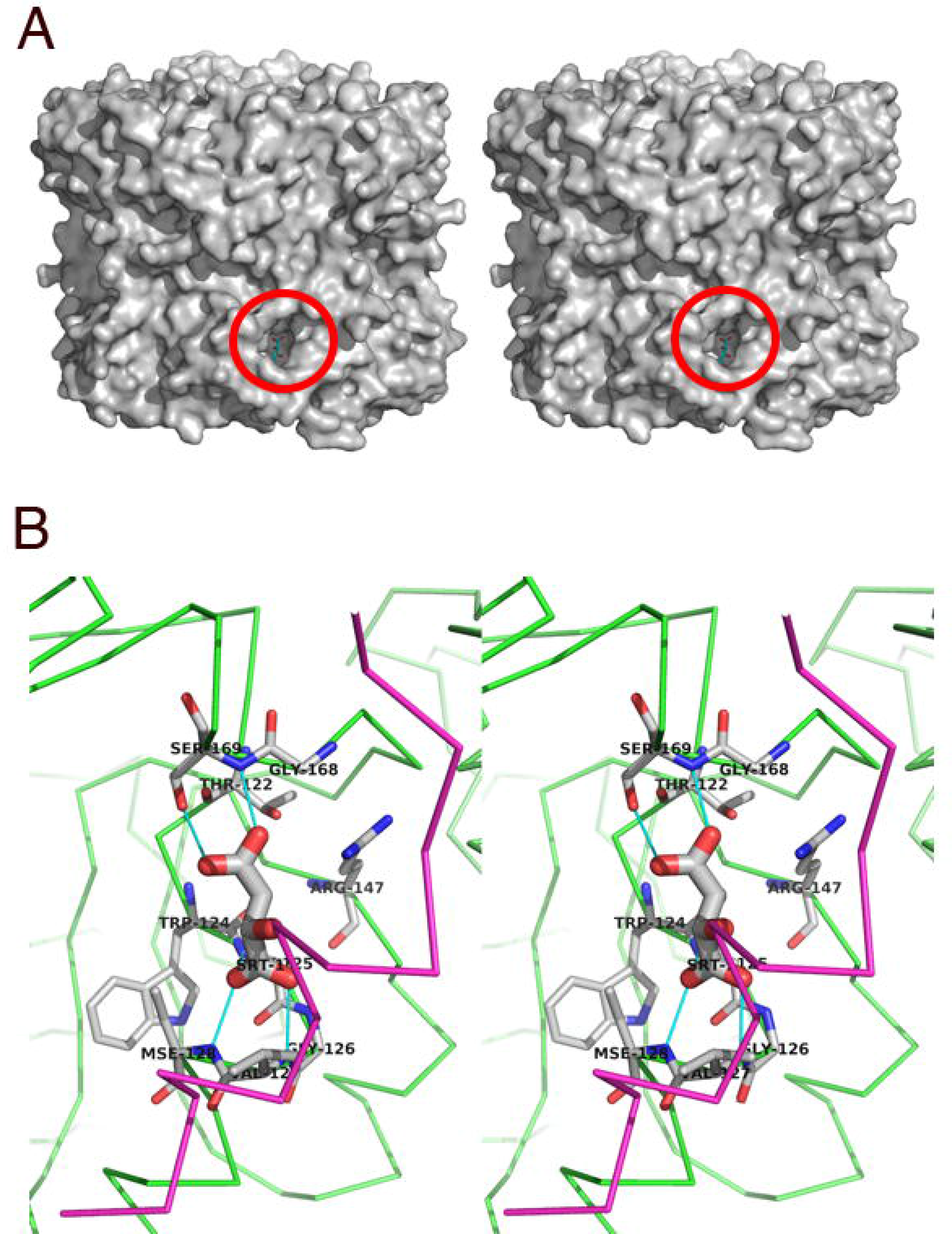
Ligand binding site of HLB5. **(A)** A stereo view of the of hexamer in solvent accessible surface representation showing a pocket with tartrate. One of the six ligand binding clefts is highlighted. Stick model of tartrate is located on surface model of hexamer. **(B)** A stereo view of the tartrate binding site of HLB5. Tartrate molecule is represented using the thick stick model, while interacting residues are represented by thin sticks, other protein chains are represented as ribbons. Hydrogen bonds are presented as cyan line. Tartrate and interacting residues are labeled.

## Conclusions

We have determined the crystal structure of HLB5, a protein previously shown to hydrolyze CMC and pretreated *Miscanthus* (Piao et al. 2014), from *P. johnsonii* DSM 18315 using single wavelength anomalous diffraction. Supported by PISA analysis and size exclusion chromatography, we have determined that the biological unit of this protein is a hexamer consisting of three dimers. By comparing HLB5 with a homologous structure in PDB, HMG/CHA aldolase from *P. putida*, we propose a ligand binding site for this cellulase. Further experiments will be required to fully understand how this novel cellulase binds and hydrolyzes its substrate.

